# Accelerated memory T cell decline and tolerogenic recall responses to SARS-CoV-2 vaccination in diabetes

**DOI:** 10.1101/2025.02.08.637262

**Authors:** Emma M Jones, Caren Sourij, Martin Stradner, Peter Schlenke, Nazanin Sereban, Othmar Moser, Rachael Quinlan, Charlotte-Eve Short, Benjamin H L Harris, Michael Fertleman, Graham P Taylor, Nick Oliver, Harald Sourij, Margarita Dominguez-Villar

## Abstract

Type 1 and type 2 diabetes are associated with increased severity and mortality from respiratory virus infections, including SARS-CoV-2. Vaccination in the general population significantly reduces the risk of severe respiratory viral infection and triggers a strong, polyfunctional and lasting T cell response in healthy individuals. However, vaccine effectiveness in people with diabetes is unclear. Here we studied the magnitude and functional characteristics of vaccine-specific CD4^+^ and CD8^+^ T cell responses to the full vaccination protocol, and the recall response after a third booster dose of SARS-CoV-2 vaccine in people with type 1 and type 2 diabetes, and compared them to those of people without diabetes. We found defects in both CD4^+^ and CD8^+^ T cell memory maintenance and the functionality of the vaccine specific T cells in people with diabetes compared to people without. In those individuals with diabetes that harbored detectable vaccine-specific T cells, they displayed an unfocused, tolerogenic phenotype characterized by increased expression of IL-13 and IL-10 in T1D and T2D compared to people without diabetes. These results have implications for vaccination strategies for people with diabetes.

---

People with type 1 (T1D) and type 2 (T2D) diabetes have a higher susceptibility to, and a more severe pathology from respiratory viral infections^1,2^. In prospective cohort studies, people with T1D and T2D have an odds ratio of 3.35 and 3.42, respectively, for severe illness after severe acute respiratory syndrome coronavirus 2 (SARS-CoV-2) infection compared to people without diabetes^2^. As such, people with diabetes are often placed in high-risk categories for vaccination prioritization strategies^3,4^. In addition, despite limited understanding of the immunological response to vaccines in people with diabetes, the same vaccination protocol against respiratory viruses as the one followed for the general population is recommended for patients with diabetes, including for Influenza and SARS-CoV-2^3–5^. There is limited information about the durability, magnitude and nature of the cellular immune response to vaccination in people with diabetes, and previous studies have reported conflicting results. Most studies on hepatitis B vaccination have demonstrated reduced immunogenicity in people with diabetes^6^, while data on other vaccines, including those against Influenza virus were mostly inconclusive^4^. Regarding SARS-CoV-2 vaccination, some evidence suggests that CD4^+^ T cell cytokine responses and neutralizing antibody titers are lower in people with T2D with higher glycemic exposure compared to those with glucose levels close to target and people without diabetes^7^, while other investigations did not observe differences in antibody titers^8^

Using SARS-CoV-2 vaccination as a model, we have dissected the cellular immune response to vaccination in people with T1D or T2D, by examining both T cell memory durability and functionality after the full vaccination protocol, and the magnitude and nature of the recall response after a booster dose of SARS-CoV-2 vaccine. Participants with diabetes displayed an impairment in the durability of the S-specific T cells after the full vaccination schedule. In those diabetes participants where S-specific T cells were detected, the response was unfocused, with the expected expression of IFNγ, TNF and IL-2, alongside a significant increase in the expression of the Th2 cytokine IL-13 in S-specific CD4^+^ and CD8^+^ T cells from both diabetes groups. Recall T cell responses to a booster dose of the SARS-CoV-2 vaccine were severely impaired in participants with diabetes compared to those without diabetes. While more than 80% of normoglycemic controls harbored S-specific CD4^+^ T cells, only 30% of participants with T1D and about 20% of those with T2D had detectable S-specific CD4^+^ T cells. In those individuals where S-specific CD4^+^ T cells were identified, a significant increase in IL-13- and IL-10-producing cells was observed, suggesting a tolerogenic vaccine-specific response. In those participants with T1D or T2D in which S-specific CD8^+^ T cells were detected, they also displayed a tolerogenic phenotype, with IL-10 being co-produced with most Tc-specific cytokines. These results suggest defects in both vaccine-specific T cell memory maintenance and alterations in the nature of the vaccine-specific T cell response in diabetes and have implications for vaccination strategies in people with diabetes.

## Results

### Vaccine-specific CD4^+^ and CD8^+^ T cell memory rapidly declines in T1D

To examine the durability and quality of vaccine-specific T cell immune memory in patients with diabetes, we analyzed peripheral blood mononuclear cells (PBMC) of participants with either T1D or T2D and control individuals without diabetes (ND). We investigated the immunogenicity of SARS-CoV-2 vaccines 1-4 months after completing the full course of vaccination (2 doses) with the aim of determining whether people with diabetes maintained a similar magnitude and functional characteristics of vaccine-specific cellular responses to people without diabetes. We used a library of overlapping peptides covering the whole SARS-CoV-2 Wuhan wild type spike protein (S) and activation-induced markers to identify vaccine-specific CD4^+^ and CD8^+^ T cells after a short stimulation of total PBMC with the peptide library (Figure 1a). Participants with T1D, but not with T2D, showed a significant decrease in the frequency of S-specific CD4^+^ T cells, identified as CD154^+^CD69^+^, compared to people without diabetes (approximately 2.7-fold decrease in T1D compared to the no diabetes group, Fig. 1b). More strikingly, the number of people without diabetes in which we could detect S-specific CD4^+^ T cells at the time point examined was significantly higher than that of T1D participants (25 out of 44 ND and 14 out of 44 T1D, p=0.031), while the T2D group showed a comparable number of participants harboring S-specific CD4^+^ T cells as ND (Fig. 1c). S-specific CD4^+^ T cells from the ND group displayed enhanced antigen experience as compared to T1D, with a higher proportion of memory T cells compared to diabetes groups and a lower proportion of naïve T cells (T_NAIVE_) compared to participants with T1D (Fig. 1d-e). Thus, those with T1D and T2D displayed a significant decrease in the frequency of effector memory cells (T_EM_) compared to ND controls, and T1D participants further displayed a significant increase in naïve S-specific CD4^+^ T cells compared to ND (Figure 1f). These results were not due to a difference in the distribution of memory CD4^+^ T cells among the three study groups, as no differences were observed in the frequency of central memory (T_CM_), T_EM_ and T_EMRA_ in total CD4^+^ T cells among groups (Supplementary Figure 1). These results suggest that in people with T1D and T2D, S-specific memory CD4^+^ T cell responses wane more rapidly than those of people without diabetes and upon antigen challenge, S-specific CD4^+^ T cells are activated mostly from the naïve T cell pool.

**Figure 1.**
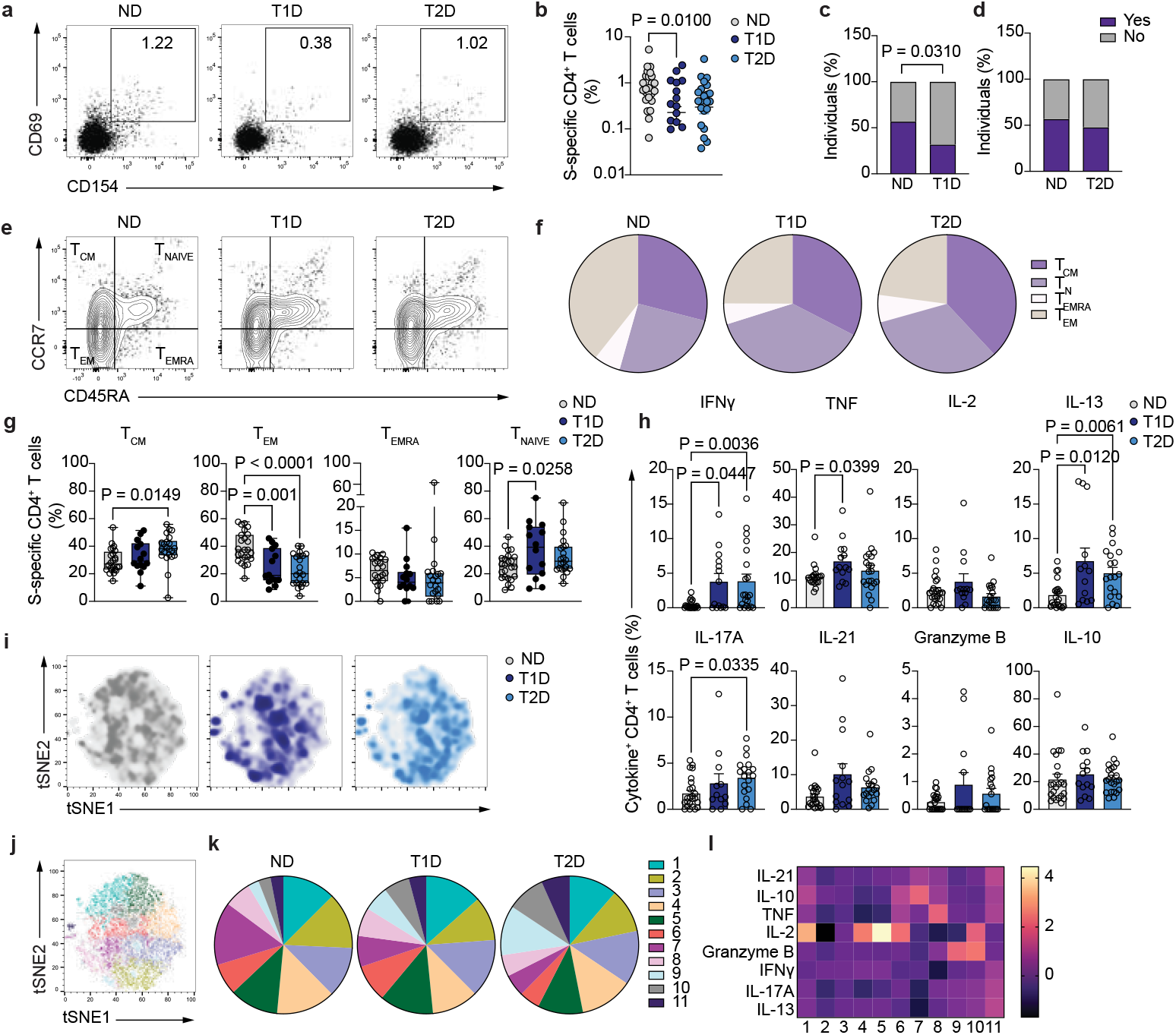
Vaccine-specific CD4^+^ T cell maintenance. **a**. Representative dot plot of S-specific CD4^+^ T cells (CD154^+^CD69^+^) after stimulation with SARS-CoV-2 spike peptide pool in people without diabetes (ND, left), T1D (middle) and T2D (right) participants. **b**. Summary of frequency of S-specific CD4^+^ T cells from total CD4^+^ T cells (n=24 ND, n=14 T1D and n=22 T2D). **c**. Percentage of individuals in whom S-specific CD4^+^ T cells were (purple) or not (grey) detected in ND controls (n=44) and T1D patients (n=44). **d**. Percentage of individuals in whom S-specific CD4^+^ T cells were (purple) or not (grey) detected in ND controls (n=44) and T2D patients (n=46). **e**. Representative dot plot of the expression of CD45RA and CCR7 in S-specific CD4^+^ T cells from ND individuals (left), T1D (middle) and T2D (right) participants. **f**. Percentage of S-specific CD4^+^ T cells from ND controls (left, n=24), T1D (middle, n=14) and T2D (right, n=22) with a naïve, memory, central memory or effector memory cells re-expressing CD45RA phenotype. **g**. Summary of S-specific CD4^+^ T cells from ND controls (n=24), T1D (n=14) and T2D (n=22) patients with a naïve, memory, central memory or effector memory cells re-expressing CD45RA phenotype. **h**. Summary of the frequency of cytokine-expressing S-specific CD4^+^ T cells from ND (n=21-24), T1D (n=14) and T2D (n=19-22) participants. **i**. tSNE projection of S-specific CD4^+^ T cells in ND (left), T1D (middle) and T2D (right) participants. **j**. tSNE projection of S-specific CD4^+^ T cell clusters using cells from the 3 study groups. **k**. Percentage of S-specific CD4^+^ T cells in each identified cluster for ND (left), T1D (middle) and T2D (right) individuals. **l**. Heatmap of geometric mean fluorescence intensity displayed as modified z-scores using median values (row-scaled). One-way ANOVA with Tukey’s correction for multiple comparisons (**b, g)**, Two-tailed Fisher’s exact test (**c, d**), Kruskal-Wallis test with correction for multiple comparisons (**h**).

The production of multiple cytokines by T cells has been associated with productive immune responses to vaccines^9–11^, so we further characterized the polyfunctionality of S-specific CD4^+^ T cells by determining the expression of cytokines associated with an optimal response to vaccination including IFNγ, TNF, IL-2^12,13^. We also included in the analysis other cytokines that are typically secreted by various helper T cell populations, such as IL-13, IL-17A, IL-21, granzyme B and the anti-inflammatory cytokine IL-10 (Figure 1g). A significantly increased proportion of S-specific CD4^+^ T cells produced IFNγ and TNF in participants with T1D compared to ND, and S-specific CD4^+^ T cells from participants with T2D displayed an increased expression of IFNγ and IL-17A. However, common to both T1D and T2D S-specific CD4^+^ T cell responses was the increased secretion of the Th2-related cytokine IL-13, suggesting a Th2-biased vaccine response. In order to better understand the heterogeneity in the functionality of S-specific CD4^+^ T cells in the study groups, we generated tSNE plots using all concatenated S-specific CD4^+^ T cells from those individuals in which they were detected. Apparent differences could be observed in the tSNE plot from ND participants compared to those of individuals with diabetes, with more similarities between the plots from T1D and T2D (Figure 1h). Unsupervised clustering analysis identified 11 different clusters of S-specific CD4^+^ T cells (Figure 1i), and the distribution of cells within such clusters did not vary significantly between ND and participants with any diabetes (Figure 1j), with only people with T2D displaying a modest decrease in the frequency of cells in Cluster 1 and a significant increase in the frequency of cells in clusters 9 and 11 compared to ND (Supplementary Figure 2). Cytokine characterization of each cluster (Figure 1k) suggested that cluster 1 contained cells with a IL-2 and IL-10 production profile, cluster 9 mainly produced granzyme B and cluster 11 was a population with a heterogeneous pattern of cytokine secretion. These results suggest that S-specific CD4^+^ T cells in people with diabetes display an unfocused cytokine phenotype with no particular enrichment of a Th2-like population, but rather a global increase in IL-13 production with no major changes in the distribution of cells within the identified clusters among study groups.

We went on to quantify and functionally characterize CD8^+^ T cell memory to vaccination. S-specific CD8^+^ T cells, identified as CD137^+^CD69^+^ (Figure 2a) were significantly reduced in frequency in participants with T1D, but not in those with T2D, compared to ND (Figure 2b, approximately 2-fold decrease in T1D compared to healthy individuals). In addition, and similarly to the CD4^+^ T cell compartment, the frequency of individuals for which S-specific CD8^+^ T cells could be detected was significantly lower in T1D compared to ND controls (22 out of 44 ND and 10 out of 44 T1D participants, p=0.0142), while no differences were observed between T2D and ND (Figure 2c). Most S-specific CD8^+^ T cells displayed a naïve or terminally differentiated T_EMRA_ phenotype, and there were no differences in the distribution of cells in naïve and memory subtypes among study groups (Supplementary Figure 3).

**Figure 2.**
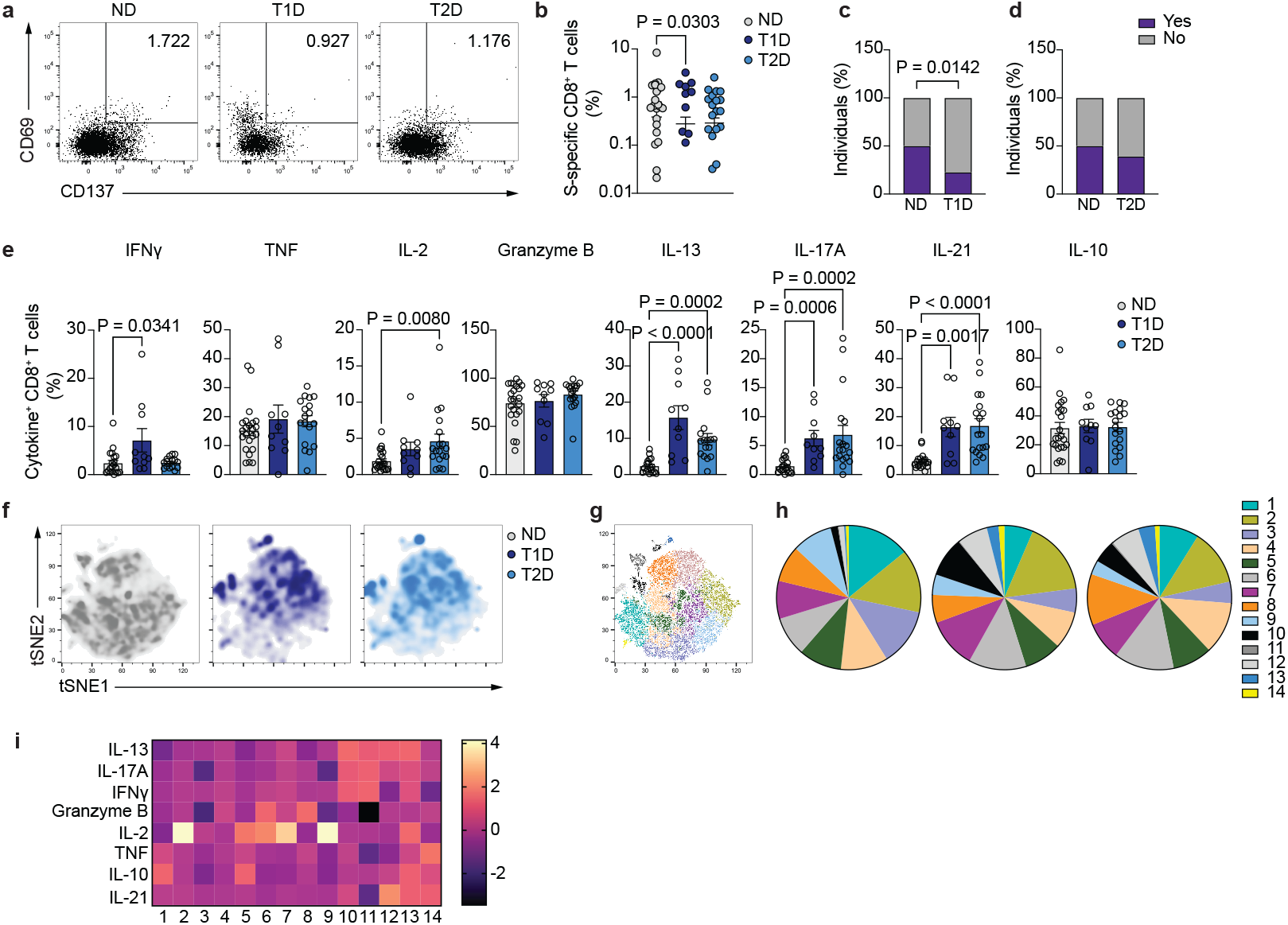
Vaccine-specific CD8^+^ T cell maintenance. **a**. Representative dot plot of S-specific CD8^+^ T cells (CD137^+^CD69^+^) after stimulation with SARS-CoV-2 spike peptide pool in ND (left), T1D (middle) and T2D (right) participants. **b**. Summary of frequency of S-specific CD8^+^ T cells from total CD8^+^ T cells (n=21 ND, n=10 T1D and n=18 T2D). **c**. Percentage of individuals in whom S-specific CD8^+^ T cells were (purple) or not (grey) detected in ND controls (n=44) and T1D participants (n=44). **d**. Percentage of individuals in whom S-specific CD8^+^ T cells were (purple) or not (grey) detected in ND controls (n=44) and T2D participants (n=46). **e**. Summary of the frequency of cytokine-expressing S-specific CD8^+^ T cells from ND individuals (n=18-22), T1D (n=10) and T2D (n=16-18) participants. **f**. tSNE projection of S-specific CD8^+^ T cells from ND controls (left), T1D (middle) and T2D (right) patients. **g**. tSNE projection of S-specific CD4^+^ T cell clusters using cells from the 3 study groups. **h**. Percentage of S-specific CD8^+^ T cells in each identified cluster for ND (left), T1D (middle) and T2D (right) participants. **i**. Heatmap of geometric mean fluorescence intensity displayed as modified z-scores using median values (row-scaled). One-way ANOVA with Tukey’s correction for multiple comparisons (**b**), two-tailed Fisher’s exact test (**c, d**), Kruskal-Wallis test with correction for multiple comparisons (**e**).

Cytokine secretion patterns in participants with diabetes were clearly different from those of ND controls (Figure 2d). Both ND and diabetes groups contained similar frequencies of TNF- and granzyme B-producing CD8^+^ T cells, with IFNγ and IL-2 production being significantly increased in T1D and T2D compared to ND, respectively. In addition, while only small frequencies of S-specific CD8^+^ T cells from the ND control group secreted other cytokines such as IL-13, IL-17A or IL-21, these were significantly increased in S-specific CD8^+^ T cells from both T1D and T2D participants, suggesting an unfocused memory response to SARS-CoV-2 vaccination. No differences were observed in the production of IL-10 among study groups (Figure 2d). Dimensionality reduction further confirmed that the global phenotype on a tSNE plot of S-specific CD8^+^ T cells in participants with diabetes was different from that of NGT (Figure 2e), and was more similar between T1D and T2D. Further corroborating the heterogeneity in the cytokine expression pattern, clustering analysis identified 14 different clusters of S-specific CD8^+^ T cells (Figure 2f). Participants with diabetes displayed an altered distribution of cells among clusters compared to ND controls, with a specific enrichment of S-specific CD8^+^ T cells in clusters 10, 12 and 13 and a significant decrease in the frequency of S-specific CD8^+^ T cells in clusters 3 and 9 (Figure 2g and Supplementary Figure 4). Clusters 10, 12 and 13 contained cells producing Th2- and Tfh-related cytokines (IL-13 in clusters 10, 11 and 12, IL-21 in clusters 12 and 13) as well as IL-17A (clusters 10 and 13) and IL-10 (cluster 13). Cluster 3 was characterized by low levels of IL-2 and IFNγ production and absence of granzyme B and IL-17A, while cluster 9 predominantly produced IL-2. These data suggest that people with diabetes display an unfocused S-specific CD8^+^ T cell memory response characterized by an increase in cells producing Th2- and Tfh-related cytokines.

### Cross-reactivity at baseline and T cell memory maintenance

Some studies have reported the presence of pre-existing T cells that recognize SARS-CoV-2 in uninfected naïve individuals, which have been suggested to enhance immune responses after SARS-CoV-2 infection and vaccination^14^. Therefore, we explored the contribution of cross-reactive T cells to the maintenance of memory to vaccination in ND controls, T1D and T2D participants (Figure 3). Correlation analyses demonstrated that the percentage of cross-reactive CD8^+^ T cells, but not of CD4^+^ T cells at baseline positively correlated with the magnitude of CD8^+^ and of both CD8^+^ and CD4^+^ S-specific T cell responses after vaccination in ND controls and T1D participants, respectively (Figure 3a). This correlation was not observed in T2D. Because there is a close association between obesity and T2D^15^, and people with obesity experience an accelerated decline in vaccine-specific B cell responses^16^, we assessed whether vaccine-specific T cell responses were correlated with obesity using body mass index (BMI) as a redout (Figure 3b). While no significant correlations were observed between S-specific CD4^+^ or CD8^+^ T cell responses after vaccination and BMI or age in ND controls or T1D group, the magnitude of S-specific CD4^+^ T cell responses was negatively correlated with BMI in the T2D group. No correlation was observed between the frequency of cross-reactive CD4^+^ or CD8^+^ T cells at baseline and BMI (data not shown). We further examined whether the magnitude of the vaccine-specific T cell response correlated with glycemia and other clinical parameters, including C-reactive protein levels and estimated glomerular filtration rate (eGFR) (Figure 3c), and only a negative correlation between the frequency of S-specific CD8^+^ T cells after vaccination and C-reactive protein levels was observed in participants with T1D.

**Figure 3.**
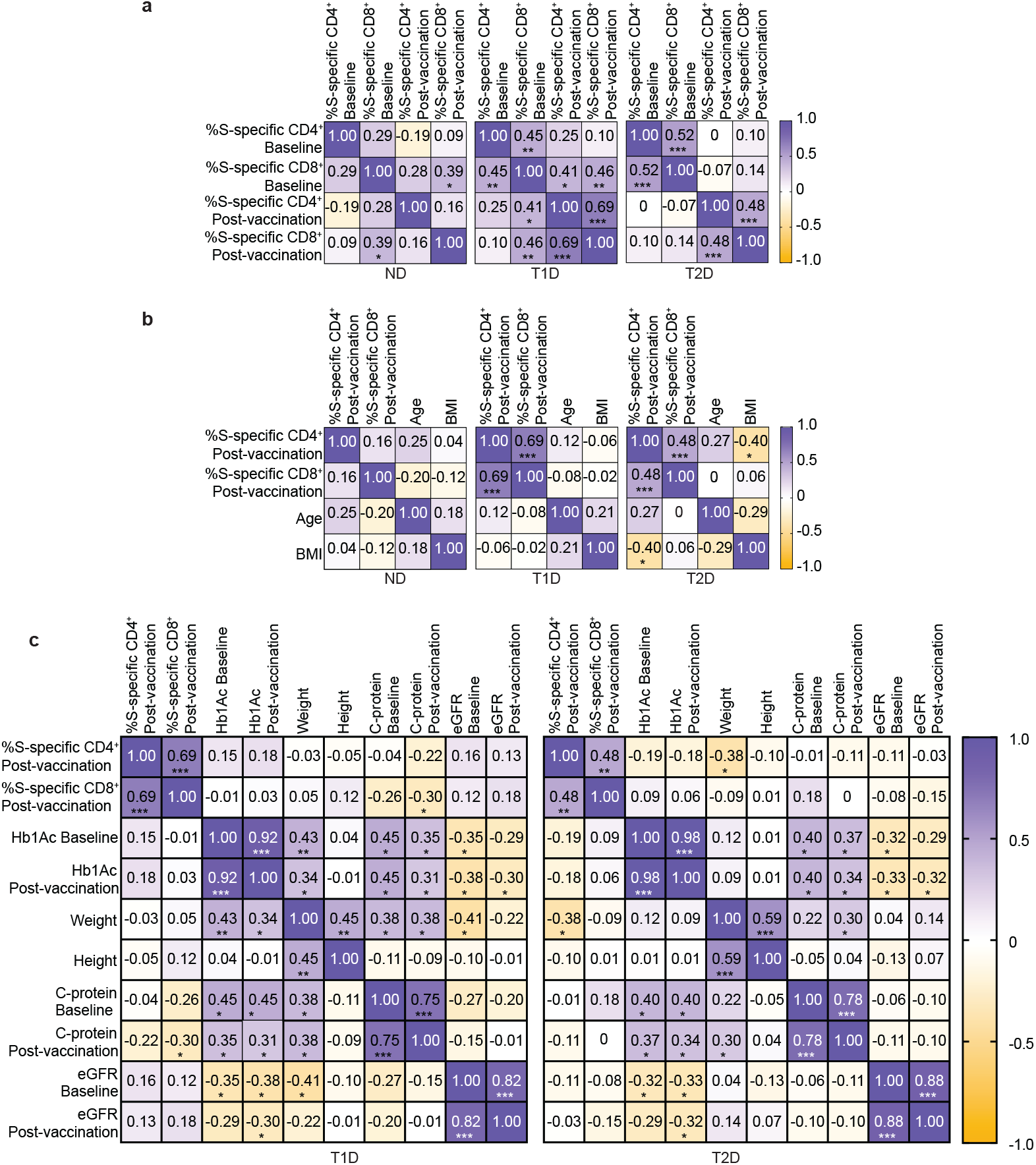
Cross-reactivity and association with clinical parameters. **a**. Correlation analysis of the frequency of S-specific CD4^+^ and CD8^+^ T cells at baseline and after full vaccination protocol in ND controls (left), T1D (middle) and T2D (right) participants. **b**. Correlation of frequency of S-specific CD4^+^ and CD8^+^ T cells after full vaccination protocol with age and body mass index (BMI) in ND controls (left), T1D (middle) and T2D (right) patients. **c**. Correlation of S-specific CD4^+^ and CD8^+^ T cell frequency with clinical parameters at baseline and after full protocol of vaccination in T1D (left) and T2D (right) participants. Spearman’s rank correlation coefficient (**a, b, c**).

These data suggest that the frequency of cross-reactive CD8^+^ T cells at baseline correlates with higher frequencies of both vaccine-specific CD4^+^ and CD8^+^ T cells after vaccination in people with T1D and ND controls, but not in people with T2D. Moreover, obesity is negatively associated with the maintenance of vaccine-specific CD4^+^ T cell memory in people with T2D.

### Anti-inflammatory CD4^+^ T cell recall responses to a booster dose of SARS-CoV-2 vaccine in participants with diabetes

We went on to examine the recall T cell response to a third booster dose of SARS-CoV-2 vaccine by characterizing the S-specific T cell responses 2-4 weeks after vaccination. The frequency of S-specific CD4^+^ T cells was significantly lower in both T1D and T2D participants when compared to that of ND controls (approximately 2- and 2.25-fold decrease in T1D and T2D, respectively, Figure 4a). Additionally, the percentage of participants with T1D or T2D from which S-specific CD4^+^ T cells were detected was significantly lower than that of ND (79% of ND compared to 33% of T1D and 22% of T2D patients, Figure 4b). Besides the decreased magnitude of the CD4^+^ T cell response, in those individuals where S-specific CD4^+^ T cells were detected, the memory phenotype distribution was different among groups (Figure 4c), and while a large proportion of ND control T cells had an effector memory phenotype, T1D T cells were preferentially naïve (Figure 4d, 4e). In addition, both T1D and T2D participants displayed a decrease in the frequency of S-specific CD4^+^ T cells with an effector memory phenotype (Figure 4e). These differences were not due to global changes in antigen experience distribution in total CD4^+^ T cells among groups (Supplementary Figure 5).

**Figure 4.**
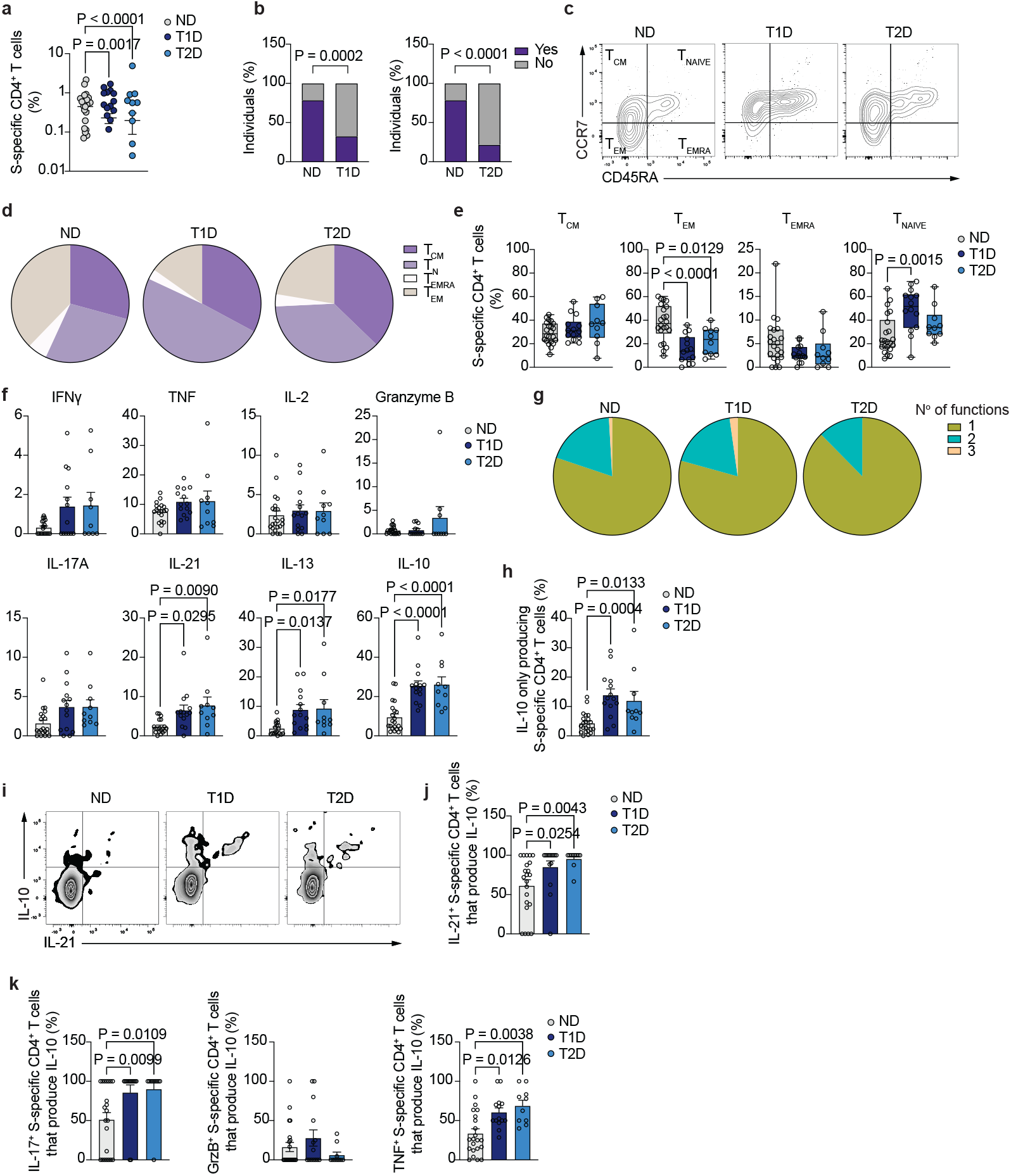
Vaccine-specific CD4^+^ T cell recall responses. **a**. Summary of frequency of S-specific CD4^+^ T cells from total CD4^+^ T cells (n=22 HC, n=14 T1D and n=10 T2D). **b**. Percentage of individuals in whom S-specific CD4^+^ T cells were (purple) or not (grey) detected in healthy individuals (n=29) and T1D patients (n=44). **c**. Percentage of individuals in whom S-specific CD4^+^ T cells were (purple) or not (grey) detected in healthy individuals (n=29) and T2D patients (n=46). **d**. Representative dot plot of the expression of CD45RA and CCR7 in S-specific CD4^+^ T cells from healthy individuals (left), T1D (middle) and T2D (right) patients. **e**. Percentage of S-specific CD4^+^ T cells from healthy individuals (left, n=22), T1D (middle, n=14) and T2D (right, n=10) with a naïve, memory, central memory or effector memory cells re-expressing CD45RA phenotype. **f**. Summary of S-specific CD4^+^ T cells from healthy individuals (n=22), T1D (n=14) and T2D (n=10) patients with a naïve, memory, central memory or effector memory S-specific CD4^+^ T cells re-expressing CD45RA phenotype. **g**. Summary of the frequency of cytokine-expressing S-specific CD4^+^ T cells from healthy individuals (n=19-22), T1D (n=12-14) and T2D (n=9-10). **h**. Distribution of S-specific CD4^+^ T cell polyfunctionality in healthy individuals (left), T1D (middle) and T2D (right) patients. **i**. Percentage of S-specific CD4^+^ T cells that only produce IL-10. **j**. IL-10 and IL-21 production by S-specific CD4^+^ T cells. **k**. Summary of the percentage of IL-21-producing, S-specific CD4^+^ T cells that co-produce IL-10. **l**. Summary of the percentage of IL-17A-(left), granzyme B-(middle) and TNF-producing (right) S-specific CD4^+^ T cells that co-produce IL-10. One-way ANOVA with Tukey’s correction for multiple comparisons (**a, f, I, k, l)**, Two-tailed Fisher’s exact test (**b, c**), Kruskal-Wallis test with correction for multiple comparisons (**g**).

The functionality of S-specific CD4^+^ T cells was assessed by dissecting the pattern of cytokine production (Figure 4f). In those individuals in which S-specific CD4^+^ T cells were detected, no significant changes in the production of IFNγ, TNF, IL-2 or granzyme B were observed among ND, T1D and T2D participants. Polyfunctionality analysis using these 4 cytokines^13^ (Figure 4g) demonstrated that the majority (>75%) of the vaccine-specific CD4^+^ T cells in the 3 groups were producers of a single cytokine (either IFNγ, TNF, IL-2 or granzyme B), and most of the remaining cells were bi-functional. However, no significant differences in the percentage of single- or polyfunctional cells were found among ND controls, T1D and T2D groups Supplementary Figure 6).

We also measured the frequency of S-specific CD4^+^ T cells producing IL-17A, IL-21, IL-13 and IL-10, and while no differences in IL-17A production were observed, S-specific CD4^+^ T cells from T1D and T2D displayed an increased frequency of IL-21-, IL-13- and IL-10-producing cells, suggesting an unfocused, Th2-biased helper T cell response to vaccination (Figure 4f). We decided to explore further the origin of the increased secretion of IL-10 to determine whether the elevated levels of IL-10 were distributed among cells producing other cytokines. The frequency of cells producing only IL-10 and no other measured cytokine was significantly higher in both T1D and T2D participants compared to ND controls (Figure 4h). Moreover, the increase in IL-21-producing CD4^+^ T cells in participants with diabetes, which are important in providing B cell help and in enhancing the survival and function of cytotoxic CD8^+^ T cells upon vaccination or viral infection, was accompanied by a concomitant increase in IL-10 production (Figure 4i). Thus, while only about 50% of IL-21-producing CD4^+^ T cells in HC were co-expressing IL-10, more than 85% and 95% S-specific, IL-21-producing CD4^+^ T cells were also producing IL-10 in T1D and T2D, respectively (Figure 4j). Additionally, IL-10 co-production was not restricted to IL-21-producing cells, and most of IL-17A-secreting S-specific CD4^+^ T cells were also expressing IL-10 in T1D and T2D, compared to approximately 50% in ND control (Figure 4k). Similarly, a significantly increased percentage of TNF-producing CD4^+^ T cells was co-producing IL-10 in T1D and T2D participants compared to ND (60% and 69% in T1D and T2D, respectively, compared to 33% in ND controls).

These data suggest that CD4^+^ T cell recall responses to SARS-CoV-2 vaccination are significantly impaired in people with diabetes. Besides the significant decrease in participants in which detectable responses are identified at the time point analyzed, detectable S-specific CD4^+^ T cells are characterized by an anti-inflammatory, IL-10-mediated phenotype that includes not only the appearance of an IL-10-producing population, but also the acquisition of IL-10 expression by most helper CD4^+^ T cells.

### Anti-inflammatory CD8^+^ T cell recall response to a third booster dose of SARS-CoV-2 vaccine in participants with diabetes

We subsequently analyzed the vaccine-specific CD8^+^ T cell recall response in ND controls, T1D and T2D participants (Figure 5). The frequency of S-specific CD8^+^ T cells after the SARS-CoV-2 vaccine booster dose was similar among ND, T1D and T2D participants, and no significant differences were observed in the percentage of individuals in whom S-specific CD8^+^ T cells were detected between ND and T1D groups (Figure 5a). However, a significant decrease in the percentage of individuals with T2D that had detectable vaccine-specific CD8^+^ T cell responses compared to ND was observed (Figure 5b). The distribution of antigen experience of vaccine-specific CD8^+^ T cells was not significantly different among ND controls and people with diabetes (Figure 5c), with similar proportions of S-specific CD8^+^ T cells with a naïve and T_EMRA_ phenotype, and lower proportions of effector memory and central memory phenotypes (Figure 5d). This cellular distribution of antigen experience was not different from that of total CD8^+^ T cells from the three study groups, although diabetes groups showed a modest increase in the frequency of total CD8^+^ T cells with a central memory phenotype (Supplementary Figure 7). To assess functionality, the secretion of TNF, IFNγ, IL-2 and granzyme B were measured as a readout for an efficient CD8^+^ T cell response to vaccination^13^. In addition, IL-21, IL-17A, IL-13 and IL-10 were assessed as readouts for cells with Tc21, Tc17, Tc2 and regulatory phenotypes, respectively (Figure 5e). Participants with T1D and with T2D harbored an increased frequency of TNF- and IL-2-secreting S-specific CD8^+^ T cells, while no differences were observed in the production of IFNγ or granzyme B compared to ND controls. We used these 4 cytokines to explore polyfunctionality of the CD8^+^ T cell recall response, and while most cells were expressing only one function, there was a significant increase in the percentage of S-specific CD8^+^ T cells that produced a combination of 2 and 3 cytokines in T2D participants compared to ND (Figure 5f). S-specific CD8^+^ T cells in T1D participants also displayed an increased frequency of cells co-producing 3 cytokines, suggesting a significant increase in polyfunctionality in both T1D and T2D patients compared to ND (Supplementary Figure 8).

**Figure 5.**
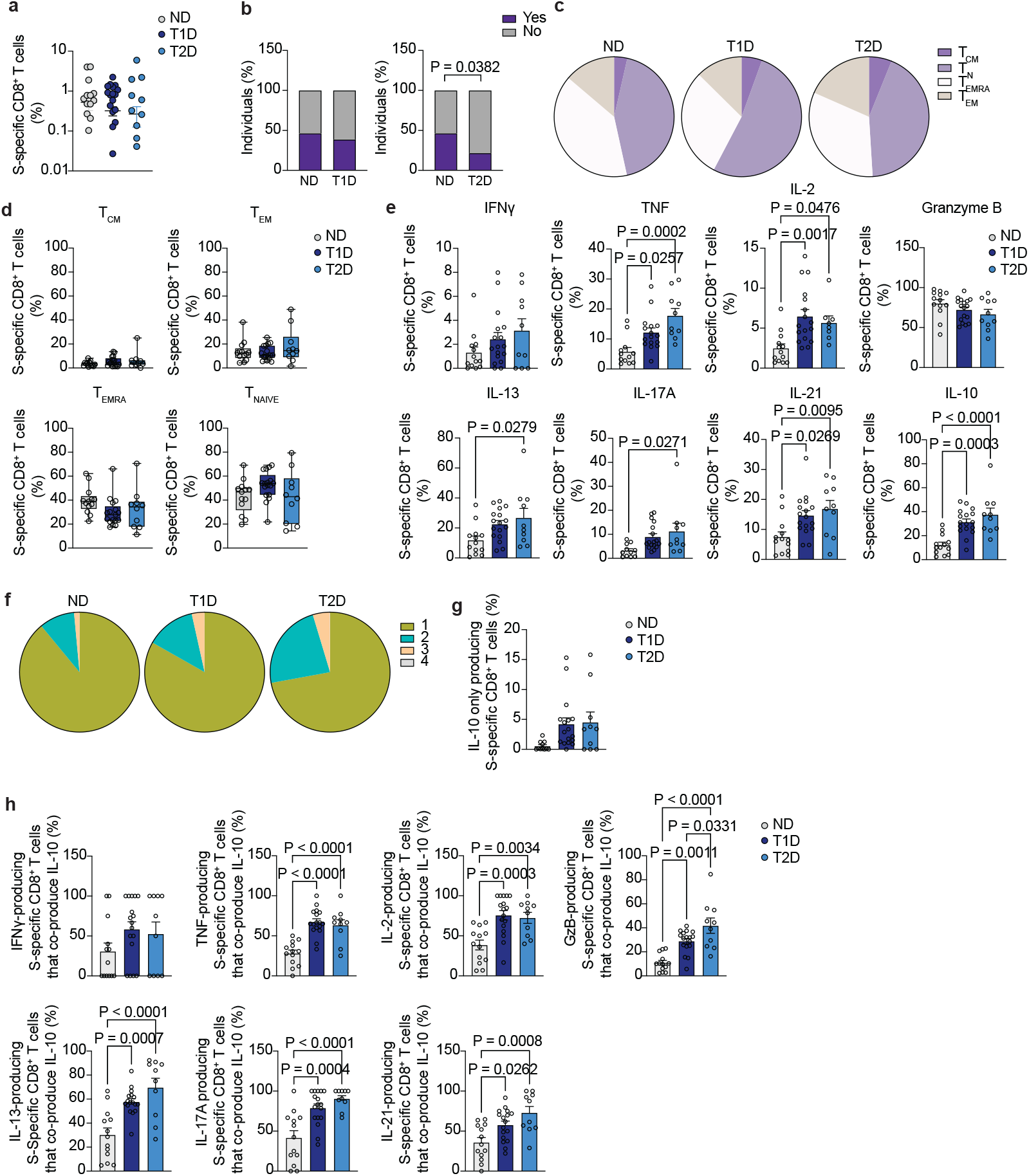
Vaccine-specific CD8^+^ T cell recall responses. **a**. Summary of frequency of S-specific CD8^+^ T cells from total CD8^+^ T cells (n=13 HC, n=17 T1D and n=10 T2D). **b**. Percentage of individuals in whom S-specific CD8^+^ T cells were (purple) or not (grey) detected in healthy individuals (n=29) and T1D patients (n=44). **c**. Percentage of individuals in whom S-specific CD8^+^ T cells were (purple) or not (grey) detected in healthy individuals (n=29) and T2D patients (n=46). **d**. Percentage of S-specific CD8^+^ T cells from healthy individuals (left, n=13), T1D (middle, n=17) and T2D (right, n=10) with a naïve, memory, central memory or effector memory cells re-expressing CD45RA phenotype. **e**. Summary of S-specific CD8^+^ T cells from healthy individuals (n=13), T1D (n=17) and T2D (n=10) patients with a naïve, memory, central memory or effector memory S-specific CD8^+^ T cells re-expressing CD45RA phenotype. **f**. Summary of the frequency of cytokine-expressing S-specific CD8^+^ T cells from healthy individuals (n=11-13), T1D (n=16-17) and T2D (n=8-10). **g**. Distribution of S-specific CD8^+^ T cell polyfunctionality in healthy individuals (left), T1D (middle) and T2D (right) patients. **h**. Summary of percentage of S-specific CD8^+^ T cells that only produce IL-10. **i**. Summary of the percentage of cytokine-producing S-specific CD8^+^ T cells that co-produce IL-10. One-way ANOVA with Tukey’s correction for multiple comparisons (**a**, e), Two-tailed Fisher’s exact test (**b, c**), Kruskal-Wallis test with correction for multiple comparisons (**f, h, i**).

However, the frequency of cells producing IFNγ, TNF and IL-2 was significantly lower than those S-specific CD8^+^ T cells producing other helper cytokines. Thus, increased expression of IL-21 and IL-10 were detected in people with T1D compared to ND (Figure 5e). Similarly, S-specific CD8^+^ T cells from participants with T2D produced significantly increased IL-13, IL-17A, IL-21 and IL-10. These data suggest that vaccine-specific CD8^+^ T cell recall responses are unfocused and characterized by the production of IL-10 in people with diabetes (31% and 37% of IL-10-producing CD8^+^ T cells in T1D and T2D participants, respectively, compared to 12% in people without diabetes).

The significantly increased production of IL-10 by S-specific CD8^+^ T cells in diabetes groups, potentially detrimental for an optimal vaccine-specific T cell response, prompted us to examine its pattern of expression further, and we explored the possibility of a CD8^+^ T cell regulatory population arising in diabetes characterized by IL-10 expression. However, no significant increase in the frequency of S-specific CD8^+^ T cells secreting only IL-10 (and not any other cytokine assessed) was observed in the diabetes groups compared to ND (Figure 5g). Instead, when we examined whether the secretion of IL-10 was associated with any specific Tc population (identified by the expression of their canonical cytokines^17^), we found that a significant proportion of TNF-, granzyme B- and IL-2-, IL-13-, IL-17A- and IL-21-secreting cells were co-producing IL-10 in people with diabetes as compared to ND participants (Figure 5h), suggesting the acquisition of IL-10 production capacity by most Tc types (Supplementary Figure 9).

## Discussion

In this work we have dissected the cellular response to vaccination in individuals with T1D and T2D, focusing on memory maintenance, magnitude and nature of the vaccine-specific CD4^+^ and CD8^+^ T cell responses. Our results suggest an impairment in memory maintenance in T1D, both in the CD4^+^ and CD8^+^ T cell compartments, which is accompanied by a decreased frequency of vaccine-specific T cells after the full vaccination protocol. We also identified significant alterations in the functionality of these cells, with an increased secretion of the Th2 related cytokine IL-13. Recall responses to a booster dose of SARS-CoV2 vaccination were also impaired in both T1D and T2D, with a significant percentage of participants with diabetes failing to mount a S-specific T cell response at the time point analyzed. Interestingly, in those participants with diabetes where a response was detected, it was dominated by the secretion of IL-10 that was present in most cytokine-producing cells, suggesting a tolerogenic, anti-inflammatory response to the vaccine.

Studies exploring the immune responses to SARS-CoV-2 vaccination in T2D have mostly measured vaccine-specific antibody titers and show conflicting results, with some works suggesting a diminished antibody response after vaccination^7,18^ and other reports demonstrating similar antibody titers between diabetes participants and ND controls^19,20^. The durability and magnitude of vaccine-specific T cell responses in our T2D cohort, however, resembled those found in people without diabetes, in line with other studies identifying similar T cell responses to SARS-CoV-2 vaccination in ND controls and people with T2D vaccinated with the BNT162b2 vaccine. Interestingly, humoral and cellular responses to whole inactivated SARS-CoV-2 vaccines have been shown to be decreased in people with T2D^21^, and vaccine type seems to differentially affect antibody responses in diabetes^22^. The impaired immunological response to SARS-CoV-2 vaccination and booser dose in people with diabetes does not seem to be specific to SARS-CoV-2, because similar results have been observed for the hepatitis B vaccine^6^. In our study, participants with T1D displayed most of the alterations in frequency and T cell memory maintenance after the full protocol of vaccination compared to T2D and ND controls. These data are in agreement with a recent study measuring T cell derived cytokines as a readout for T cell responses to SARS-CoV-2 vaccination^23^, and an early study examining T cell responses to other vaccines in T1D^24^. Indeed, an impairment in the immune response following vaccination in patients with T1D has been previously suggested for other immunization strategies, including Influenza, rotavirus and Influenza type B^24,25^, although underlying mechanisms remained elusive. In our study, people with T1D displayed a significantly lower frequency of CD4^+^ T cells after the full vaccination protocol and after the booster dose, and this was accompanied by a significant reduction in the percentage of people with T1D with S-specific CD4^+^ T cell responses compared to ND. Several mechanisms could explain these results, including a defect in maintenance of memory CD4^+^ T cells to vaccination in people with T1D, and a delayed T cell response to vaccination (Figures 4 and 5). Due to the limited number of time points available for analysis after the full course of vaccination or the recall response, our results can only suggest a delayed response to vaccination as the underlying mechanism for the small number of people with T1D and T2D that responded to the booster dose at the time point analyzed.

Systems biology and clinical studies on the response to SARS-CoV-2 vaccines in humans indicate that efficient vaccine-specific CD4^+^ T cell responses are characterized by a Th1-like profile and either low or absent levels of the Th2 cytokines IL-4 and IL-5^26,27^. The unfocused and tolerogenic recall response to SARS-CoV-2 vaccination dominated by the acquisition of IL-10 and IL-13 production by vaccine-specific T cells from people with diabetes, is in contrast with the focused, TNF-, IL-2-IFNγ-producing T cell response observed by us and others^28^ in people without diabetes, where vaccination barely induced any Th2- or Th17-type cytokine. The fact that IL-10 production is acquired by CD4^+^ and CD8^+^ T cells with a variety of phenotypes and is not restricted to a specific T helper or Tc subpopulation suggest that in diabetes, vaccination generates a generalized anti-inflammatory response, potentially reducing the efficacy of the vaccine. Indeed, early induction of IL-10 limits antigen-specific CD4^+^ T cell expansion, function and secondary recall responses during phagosomal infections^29^ and IL-10 receptor blockade delivered simultaneously with BCG vaccine stimulates immunity and sustains long-term protection in mice^30^. Further investigations are warranted to unambiguously define the involvement of IL-10 and IL-13 in the maintenance and function of vaccine-specific T cell responses and their favorable or detrimental role in antiviral immunity upon infection in people with diabetes. However, existing data indicate that expansion of Th1 CD4^+^ cells with the capacity to secrete IFNγ, TNF and IL-2 upon SARS-CoV-2 and other respiratory viral infections are associated with improved protection and viral control as opposed to an enhanced type 2 response^31–35^. Therefore, our data suggest that people with T1D, and to a lesser extent, people with T2D, mount an inefficient cellular response to vaccination.

Our results have implications for SARS-CoV-2, and potentially other, vaccination regimens for people with diabetes and warrant further investigations to pinpoint the mechanisms underlying the defects observed and broader studies focused on vaccine effectiveness in diabetes populations.

### Tables

**Table 1. Participant characteristics.**

## Online Methods

### Participants and clinical data collection

The cohorts analyzed in this study consisted of individuals aged 18 years or older either without known diabetes or with a diagnosis of T1D or T2D. Diabetes participants were recruited to the COVAC-DM study, a prospective, multi-center, cohort study aimed at examining the immune response to COVID-19 vaccination in people with type 1 and 2 diabetes mellitus and a glycated hemoglobin level ≤58 mmol/mol (7.5%) or >58 mmol/mol respectively, (EudraCT2021-001459-15)^19,20^. Participants were administered the full 2-dose course and a third “booster” dose of BTN162b2 (Pfizer/BioNTech), ChAdOx1-S (AstraZeneca) (only for the first 2 vaccination doses and only used in 8 participants (5.3% of the entire study)) or mRNA-1273 (Moderna) vaccines between April 2021 and March 2022. Blood samples were collected at baseline (60-2 days before their first vaccination), 4-16 weeks after the second dose and 2-4 weeks after the third dose after informed consent was obtained. None of the T1D and T2D had had COVID-19 either before or during the length of the study. The control group without diabetes consisted in individuals from the CoVVac (EudraCT: 2021-001040-10) study and additional participants from the COVIDITY study, a prospective observational serial sampling study of whole blood that recruited people with suspected or confirmed SARS-CoV-2 infection that were followed for 18 months. 12 healthy individuals that were included in the data analysis in Figures 1 and 2 were participants from the COVIDITY study and had had COVID-19 at least 12 months before COVID-19 vaccination (Table 1). Ethical approvals were obtained from XXXXX and the South Central Oxford C Research Ethics Committee (IRAS ID 15/sc/0089).

Participants of both cohorts were recruited at the University Hospital Graz, either from established cohorts (Graz Diabetes Registry for Biomarker Research), the outpatient clinic for diabetes, lipid disorders and metabolism or via advertisements. Main exclusion criteria for the COVAC-DM trial were: active malignancy (excluding intraepithelial neoplasia of the prostate gland and the gastrointestinal tract and basalioma), pregnancy, acute inflammatory disease, immunosuppressant therapy, alcohol abuse (more than 15 standard drinks a week) or any contraindication to the vaccine as well as previous episode of COVID-19. The trial was approved by the ethics committee of the Medical University of Graz (33-366 ex 20/21) The CoVVac trial included healthy controls without known diabetes and excluded people with the presence of disease or therapy potentially interfering with the response to vaccination.

### PBMC isolation and storage

Blood samples (40 ml) will be collected using the BD Vaccutainer CPT™ (Becton Dickinson, NJ, USA) tubes to separate mononuclear cells. Peripheral blood mononucleated cells will be frozen in DMSO-based media and stored in liquid nitrogen until further analysis at the Center for Biomarker Research in Medicine (CBmed).

### Identification of S-specific T cells

PBMC were thawed in a pre-warmed water bath at 37 °C and immediately transferred to a 15 mL conical tube where 5 mL of pre-warmed (37 °C) cell culture media supplemented with 50 U/mL of Pierce™ Universal Nuclease (Thermo Fisher Scientific) or Benzonase^®^ (Merck) was added drop by drop. Cell culture media used was RPMI 1640 (GIBCO) supplemented with 2 mM L-Glutamine, 5% human serum AB (Sigma-Aldrich) and 100 µg/mL penicillin and streptomycin (GIBCO). Samples were subsequently centrifuged for 10 minutes at 300 x *g* and resuspended in warm cell culture media at 2 x 10^7^ cells/mL. The PBMCs were rested for 2 hours at 37 °C, 5% CO_2_ before *ex vivo* stimulation. 1-2 million PBMCs were incubated with 1 µg/mL anti-CD40 (Miltenyi Biotec) and 1 µg/mL anti-CD28 (Miltenyi Biotec) for 20 minutes at 37°C. Cells were then stimulated with either PepTivator^®^ SARS-CoV-2 complete S protein (Wuhan wild-type, Miltenyi Biotec) or vehicle for 16 hours. PMA (50 ng/mL) and ionomycin (250 ng/mL) stimulation was carried out in some samples as a positive control for stimulation following the same protocol. For intracellular stainings, GolgiStop (BD Biosciences) and GolgiPlug (BD Biosciences) were added to the cultures after 12 hours of stimulation for 4 hours

### Cellular staining for flow cytometry

Cells were stained with LIVE/DEAD Fixable Blue Dead Cell Dye (Thermo Fisher Scientific) according to the manufacturer’s specifications. A FcR receptor blocking step with FcR Blocking Reagent Human (Miltenyi Biotec) before cell surface antibody staining, which included antibodies against CD3 (BV785, clone UCHT1), CD4 (PerCP, clone RPA-T4), CD154 (PE, clone 24-31), CD137 (BV711, clone 4B4-1), CD69 (Alexa Fluor 700, clone FN50), CD45RA (BV650, clone HI100), CCR7 (APC-Cy7, clone G043H7), all from Biolegend. The cells were subsequently fixed using the FoxP3/ Transcription Factor Staining Buffer kit (Thermo Fisher Scientific) following the manufacturer’s specifications. The cells were then stained with intracellular antibodies, including anti-human IL-13 (BV421, clone JES10-5A2), IL-17A (BV510, clone BL168), granzyme B (FITC, clone QA16A02), IL-2 (PeCP-Cy5.5, clone MQ1-17H12), TNF (PE Dazzle 594, clone Mab11), IL-10 (PE-Cy7, clone JES3-9D7), from Biolegend and IL-21 (eFluor 660, clone eBio3A3-N2) and IFNγ (BV605, clone XMG1.2), from Thermo Fisher Scientific, washed and resuspended in 220µl of PBS.

The samples were run on a Fortessa instrument (BD Biosciences) and analyzed using FlowJo v10.0 (BD Biosciences).

### Dimensionality reduction and cluster analysis

Dimensionality reduction and tSNE plots were obtained as previously described^36,37^. Briefly, S-specific CD4^+^ T cells (or S-specific CD8^+^ T cells) from healthy individuals and patients with T1D and T2D were concatenated. The concatenated sample was used to calculate tSNE axes using 1,500 iterations, perplexity of 30 and the default learning rate. In order to obtain cell clusters, we used Phenograph plugin in FlowJo, with all compensated parameters.

### Statistical analysis

Frequency of S-specific CD4^+^ and CD8^+^ T cells was calculated by subtracting the value of the vehicle-stimulated cells for each study participant. Only those participants in which S-specific CD4^+^ or CD8^+^ T cells were detected after substraction of background were included in further analyses, i.e. cytokine analysis, antigen experience. Data were analyzed using GraphPad Prism version 10.0. Normal distribution of the data was tested using the D’Agostino and Pearson and Anderson-Darling normality tests or Shapiro-Wilk test for those datasets with a small number of data points. Normally distributed data by at least one of the two tests were analyzed using one- or two-way ANOVA when comparing more than two groups of one or two independent variables, respectively. A two-tailed t-test was used to compare two groups. For non-gaussian distributed data, Mann-Whitney U test or Kruskall-Wallis test with correction for multiple comparisons was used to compare two or more groups, respectively. Fisher’s exact test was used to determine the presence of a non-random association between two categorical variables, and Spearman’s rank correlation coefficient was used to determine relationships between two variables. Data are represented as mean ± s.e.m. Where the data are presented as box and whiskers, the boxes extend from the 25th to the 75^th^ percentile and the whiskers are drawn down to the minimum and up to the maximum values. Horizontal lines within the boxes denote the media. P values <0.05 were considered statistically significant.

## Supporting information

Supplementary Figue 3

## Data availability

The raw numbers for charts and graphs are available in the Source data file whenever possible.

## Acknowledgements

We thank the participants who volunteered for this study and the clinical teams of the COVAC-DM, CoVVac and COVIDITY studies for patient recruitment and blood collection. EJ is a Diabetes UK PhD scholar. This work was funded by Diabetes UK (MDV). The COVAC-DM study was funded by the Austrian Science Fund (KLI-1076 to HS). We would like to thank Christine Schwarz who was involved in the early stages of the COVAC-DM study design and Barbara Prietl and Faisal Aziz who helped with the sample processing and Susanne Kaser, who collaborated in the COVAC-DM trial.

## Author contribution

EMJ performed experiments, analyzed data and wrote the manuscript. CS, MS, PS, HS, OM and MSer designed and performed the COVAC-DM and CoVVac trials. HS, MS and CS obtained funding for the clinical trials. CES and GPT designed, supervised and managed the COVIDITY observational study, RQ recruited and processed samples from the COVIDITY study, BHLH and MF recruited participants for COVIDITY study. NO designed the study and wrote the manuscript. MDV designed the study, analyzed data, wrote the manuscript and obtained funding. All authors revised and contributed to the editing of the manuscript.

## Extended data figures

**Supplementary Figure 1. Total CD4**^**+**^ **T cell antigen experience. a**. Representative dot plot of the expression of CD45RA and CCR7 in ND controls (left), T1D (middle) and T2D (right) participants. **b**. Percentage of CD4^+^ T cells from ND (left), T1D (middle) and T2D (right) individuals with a naïve, memory, central memory or effector memory cells re-expressing CD45RA phenotype.

**Supplementary Figure 2. S-specific CD4**^**+**^ **T cell cluster distribution**. Percentage of S-specific CD4^+^ T cells in each of the identified clusters. One-way ANOVA with Tukey’s correction for multiple comparisons.

**Supplementary Figure 3. Antigen experience of S-specific (a) and total (b) CD8**^**+**^ **T cells**.

**Supplementary Figure 4. S-specific CD8**^**+**^ **T cell cluster distribution**. Percentage of S-specific CD8^+^ T cells in each of the identified clusters. One-way ANOVA with Tukey’s correction for multiple comparisons.

**Supplementary Figure 5. Antigen experience in CD4**^**+**^ **T cell recall responses**. Summary of percentage of total CD4^+^ T cells from ND controls, T1D and T2D participants with a naïve, memory, central memory or effector memory cells re-expressing CD45RA phenotype. One-way ANOVA with Tukey’s correction for multiple comparisons.

**Supplementary Figure 6. Polyfunctionality of S-specific CD4**^**+**^ **T cells**. Percentage of S-specific CD4^+^ T cells from ND controls, T1D and T2D participants that co-express one, two, three or four cytokines (functions). One-way ANOVA with Tukey’s correction for multiple comparisons.

**Supplementary Figure 7. Antigen experience in CD8**^**+**^ **T cell recall responses**. Summary of percentage of total CD8^+^ T cell from ND, T1D, and T2D participants with a naïve, memory, central memory or effector memory cells re-expressing CD45RA phenotype. One-way ANOVA with Tukey’s correction for multiple comparisons.

**Supplementary Figure 8. Polyfunctionality of S-specific CD8**^**+**^ **T cells**. Percentage of S-specific CD8^+^ T cells from ND controls, T1D and T2D participants that co-express one, two, three or four cytokines (functions). One-way ANOVA with Tukey’s correction for multiple comparisons.

**Supplementary Figure 9. Co-expression of IL-10 by S-specific, cytokine-producing CD8**^**+**^ **T cells**. Dot plots show the co-expression of IL-10 and other cytokines in S-specific CD8^+^ T cells from ND (left), T1D (middle) and T2D (right) individuals.

