## Supplementary figures and images for "Accelerated memory T cell decline and tolerogenic recall responses to SARS-CoV-2 vaccination in diabetes"

### Supplementary Figue 3

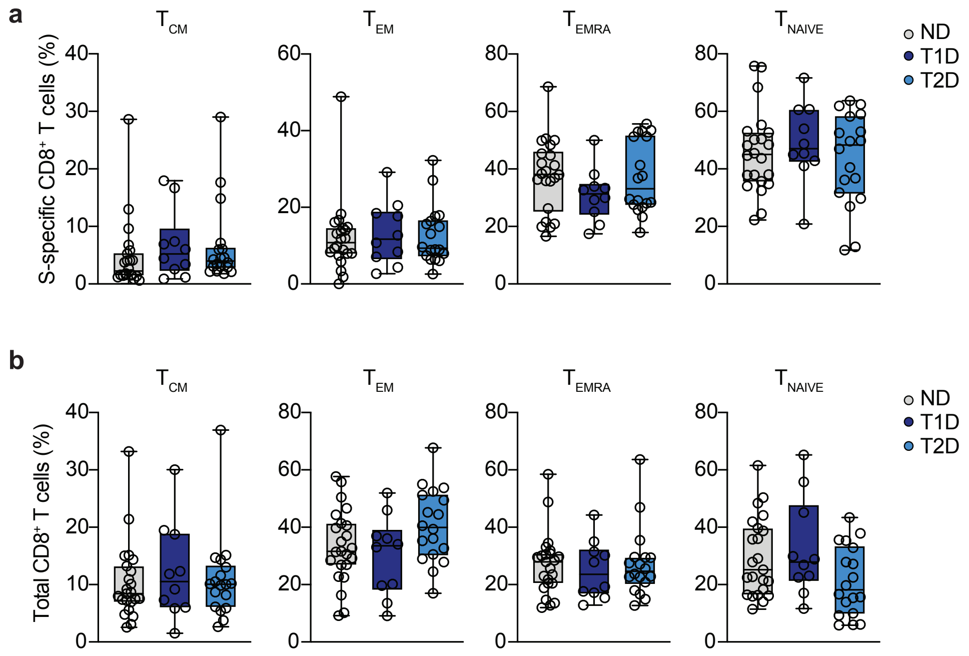


Supplementary Figure 3

### Supplementary Figure 1

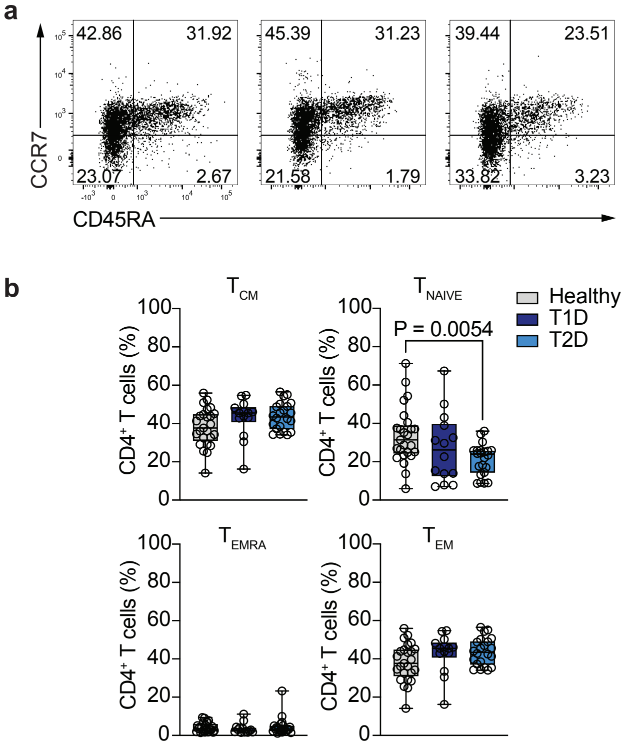


Supplementary Figure 2

### Supplementary Figure 2

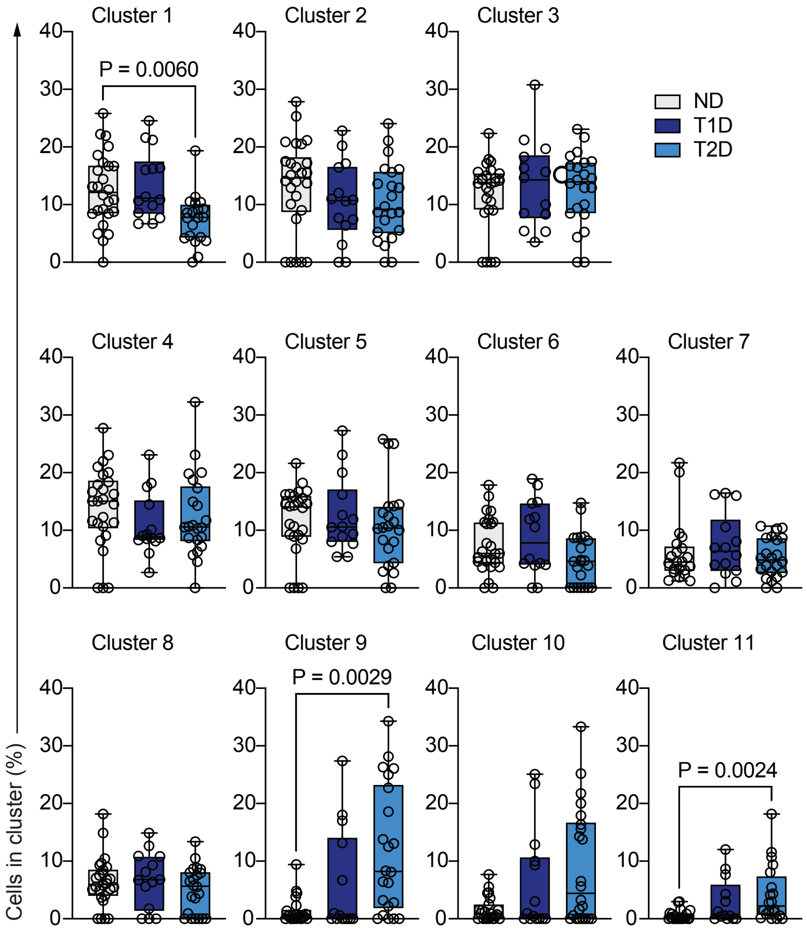


Supplementary Figure 2

### Supplementary Figure 4

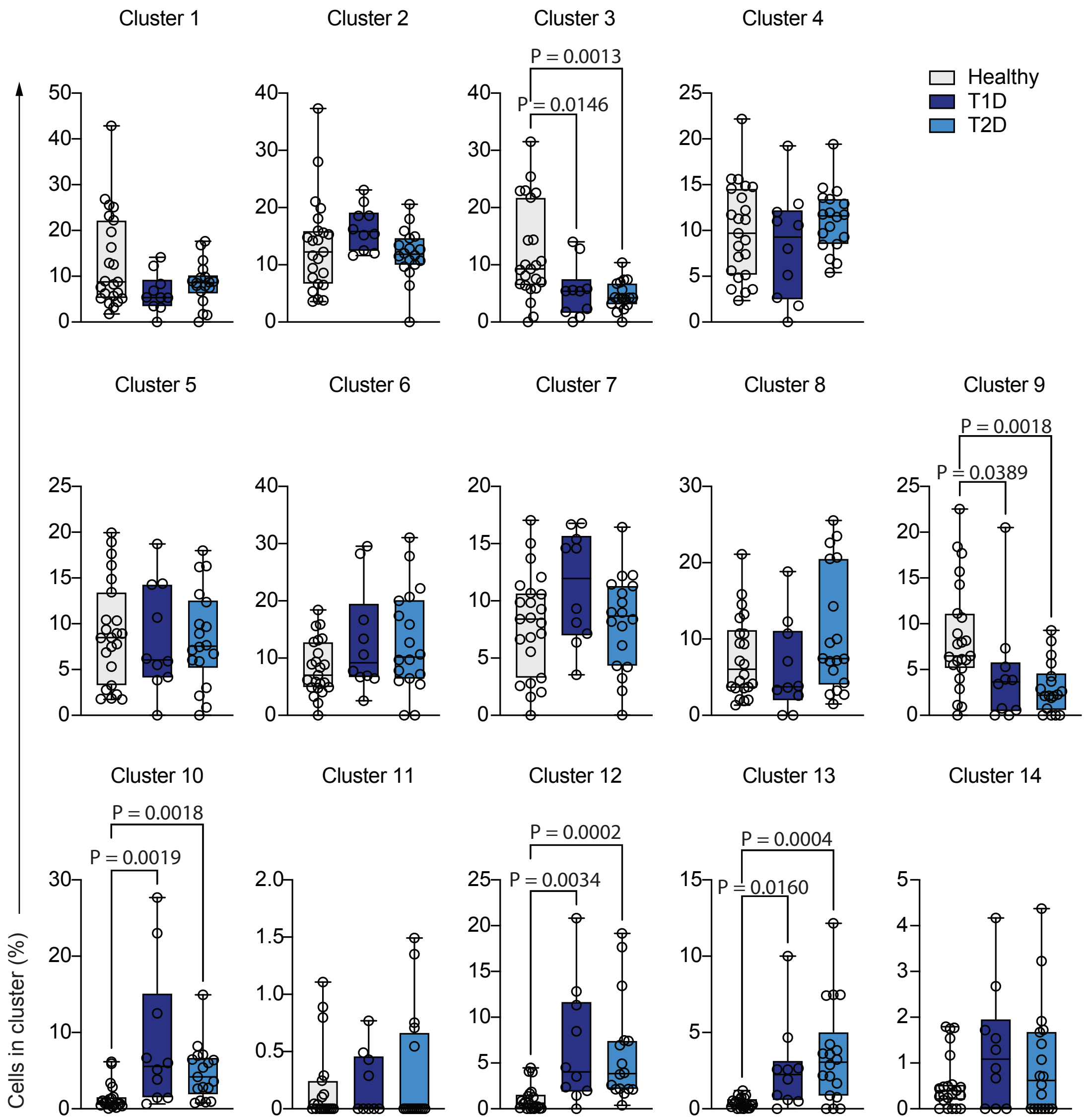


Supplementary Figure 5

### Supplementary Figure 5

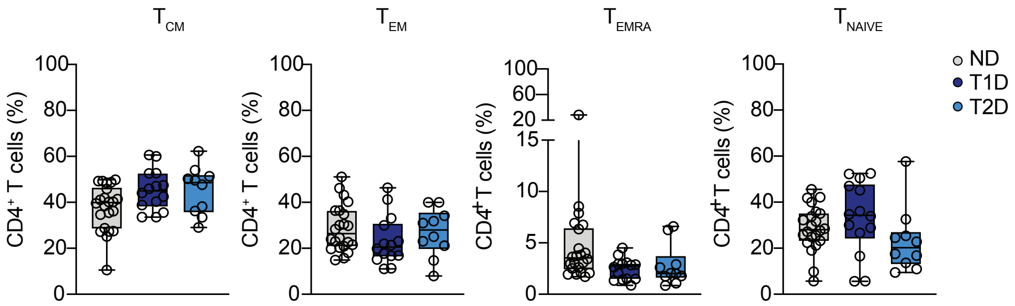


Supplementary Figure 5

### Supplementary Figure 6

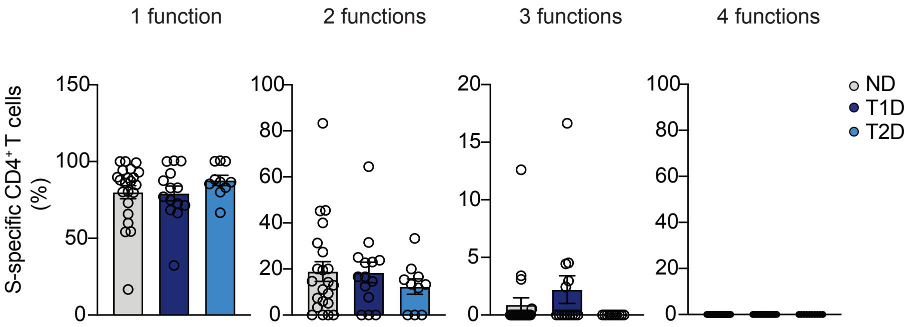


Supplementary Figure 6

### Supplementary Figure 7

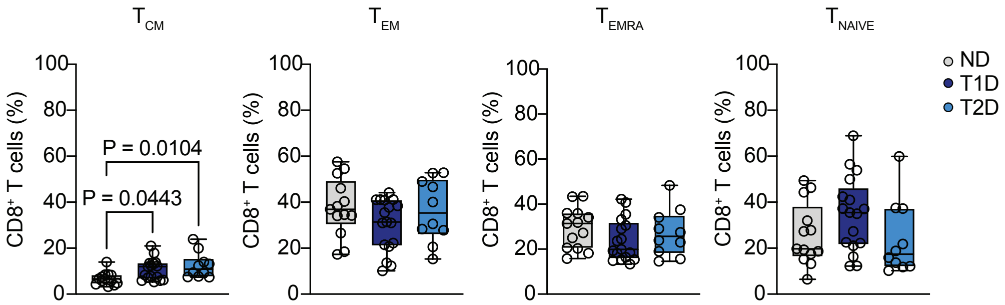


Supplementary Figure 7

### Supplementary Figure 8

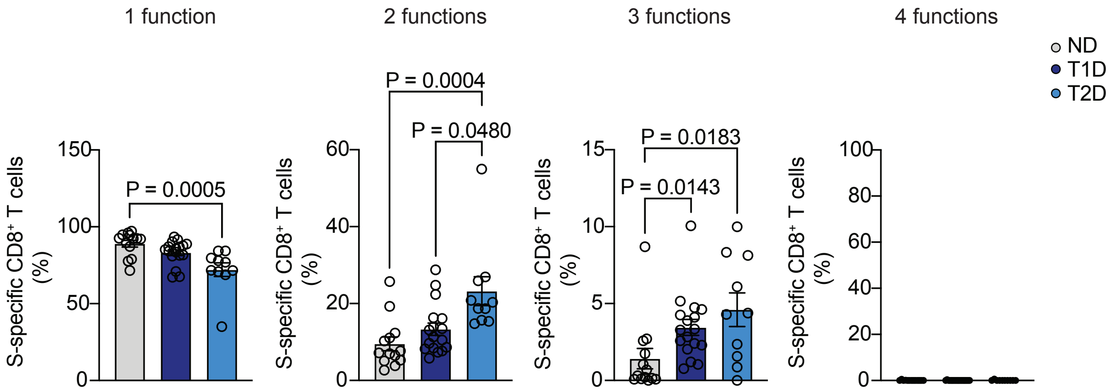


Supplementary Figure 8

### Supplementary Figure 9

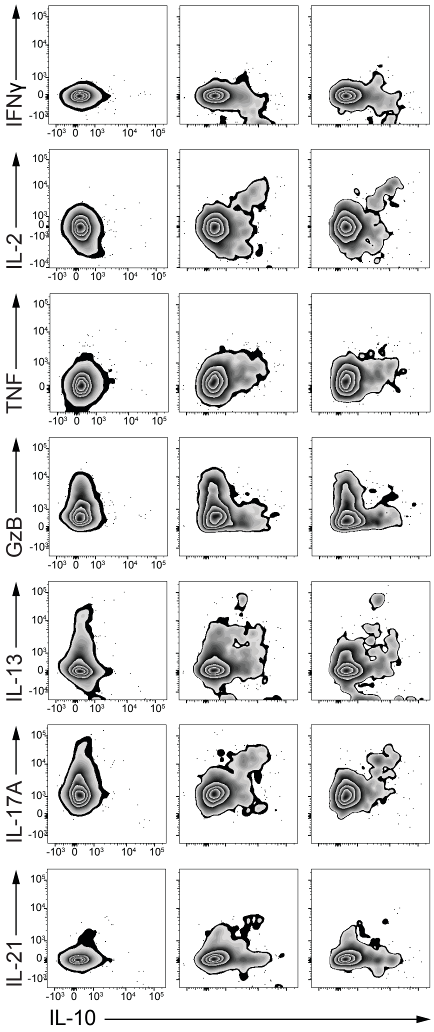


Supplementary Figure 11
